# Long-term conservation effects of protected areas in stochastic population dynamics

**DOI:** 10.1101/2021.02.06.430090

**Authors:** Nao Takashina

**Affiliations:** Department of International Studies, The University of Tokyo, 5-1-5 Kashiwa, Chiba 277-0459, Japan

**Keywords:** Ecosystem management, marine protected areas, population fluctuation, protected areas, stochastic differential equations, stochastic processes

## Abstract

Terrestrial and marine protected areas are essential tools in mitigating anthropogenic impacts and promoting population persistence and resource sustainability. Adequately implemented protected areas (PAs) promote long-term conservation effects. Stochasticity causes fluctuations in the conservation effects of PAs, and so it is important to investigate the variabilities of these conservation effects to inform their long-term conservation effects. To investigate long-term conservation effects, I develop and analyze new models of stochastic processes that encompass the fluctuations generated by demographic or environmental stochasticity in PAs management. The stochastic model is built upon individual processes. In the model, density-independent mortality, migration between PAs and non-PAs, and site preferences characterize the features of the PA. The effect of PAs size is also examined. The long-term conservation effects are quantified using the coefficient of variation (CV) of population size in PAs, where a lower CV indicates higher robustness in stochastic variations. Typically, the results from this study demonstrate that sufficiently reduced density-independent mortality in PAs and high site preference and immigration rate of PA are likely to decrease the CV. However, different types of stochasticity induce rather different consequences: under demographic stochasticity, the CV is always reduced when PAs increase the population size therein, but an increased population size by PAs does not always decrease the CV under environmental stochasticity. The deterministic dynamics of the model are investigated, facilitating effective management decisions.

## 1 Introduction

Terrestrial and marine protected areas are being expanded worldwide in response to increasing concern about species loss [45]. These protected areas (PAs) have become essential tools for mitigating anthropogenic impacts, promoting population persistence and resource sustainability, and enhancing ecological resilience [22, 35, 44, 45]. The establishment of PAs is not in itself a goal, but PAs are assumed to promote long-term conservation [28]. Existing strategic PA site selection methods face difficulties in envisioning their long-term conservation effects, because site selection often involves a snapshot of optimality, rather than a long-term consideration of the optimum [33].

In the literature, several long-term benefits of PAs arise in deterministic models under the assumption that populations approach an equilibrium state after the PAs are implemented, such as improvement of yields [39] and mitigating bycatch [18] in fisheries management. This concept is ubiquitous and characterizes an expected long-term effect of conservation practice and management (e.g., [6, 19]). In nature, however, population size fluctuates according to demographic and/or environmental stochasticity, with the former attributed to the probabilistic nature of (intrinsic) demographic events and the latter attributed to the noise induced by external factors [24, 32]. Consequently, ecological status (and, hence, PA effects) fluctuates over time (Fig. 1a), perhaps in different ways under different types of stochasticity. In such situations, effects of PAs are no longer persistent, and a population may undergo a period during which a given conservation effect is weak, potentially enhancing the risk of extinction. Therefore, variations in conservation effects, along with their average effects, are critically important to understand effects of PAs. However, prevailing equilibrium discussions cannot deal with such variations in conservation effects. Nonequilibrium discussions, however, can inform long-term PA effects (Fig. 1b), and are the focus of this study.

**Figure 1:**
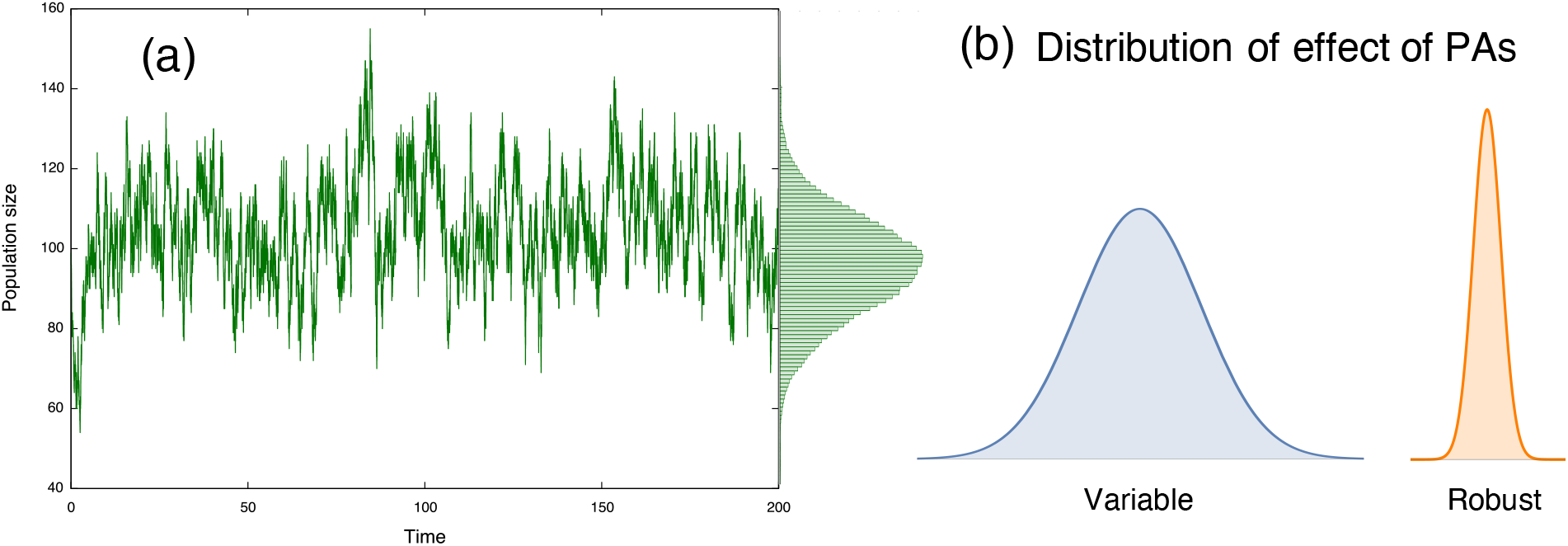
(a) Population size in protected areas (PAs) show variations over time; and (b) such variations characterizes the long-term effect of PAs. A large variation indicates that the effect is variable over time, while a small variation suggests a robust long-term conservation effect.

In practice, long-term field observations are very costly; hence, there are limited time-series data available [14]. Instead, modeling has power to effectively explore such long-term effects. Most modeling frameworks integrating stochastic events based on PA effects are in the context of marine protected areas (MPAs) (e.g., [3, 20, 26]), which are often associated with fisheries management and focus on optimal harvesting strategies [7], tradeoffs between conservation effects and fisheries profits [9,15,27], population persistence [3,47], and resilience [1, 5, 46]. Few studies have investigated the stochastic influences on long-term MPA effects in a fisheries context, or how MPAs affect variability in population size or catch [12, 15, 26]. Barnett and Baskett [5] demonstrated that, by assuming stochastic recruitment in predatory fish species, the coefficient of variation in the catch could be reduced in the case of fisheries management using MPAs. In a study of metapopulation dynamics in the Great Barrier Reef Marine Park, Hopf et al. [20] demonstrated that marine reserves could promote the stability of populations and fishery yields, regardless of fishing intensity. However, Hopf et al. [20] also showed that this conclusion depends on the location of reserves: more variable biomass can occur if disturbed reefs are protected, compared to protecting undisturbed sites. This suggests a need to understand the underlying mechanisms in determining of conservation effects. The models used in these studies target marine species and often have complex structures to describe the life histories of species. Hence, our understanding of stochasticity and PA effects is very limited, despite their broad applicability to terrestrial and marine ecosystems.

This paper develops general theoretical insights regarding the long-term effects of stochasticity on PAs (and MPAs), without restricting the discussion to fisheries management or marine environments. I focus on (i) whether PAs suppress stochastic fluctuations (i.e., providing a long-term conservation effect) and under what conditions, if any, this is achieved; and (ii) whether demographic and environmental stochasticity affect the long-term conservation effects of PAs in different ways. The developed master equation allows existing analytical methods to be used, and I develop analytical insights into the stochastic population model. This would be a difficult task using the existing complex models. This is not merely for mathematical understanding, but provides a more explicit underlying mechanism of variability in the effects and parameter dependence of PAs.

In this paper, I address these questions using a spatially explicit stochastic population model (e.g., [13, 16, 30]). I begin by looking at individual processes and formulating a master equation, and then obtain the corresponding stochastic differential equations (SDEs). While the derived stochastic population models with PAs are new, it turns out that they are stochastic analogous of existing deterministic models used to analyze equilibrium properties in MPA management. I measure the time variations of PA effects via the coefficient of variation (CV), where a smaller value indicates a more robust long-term conservation effect against stochasticity, and *vice versa*. I use the CV to quantify time-variable PA effects, which are scaled by the mean population size (note that, in general, the variance increases with the mean), but I present the CV along with the mean value.

My approach enables the long-term conservation effect of PAs under stochasticity to be discussed. Multiple parameters determine the quality of PAs, and I can examine various biological and management scenarios. Additionally, by defining “inappropriate” PAs as sites with a higher mortality than non-PAs (e.g., caused by illegal use of protected species within PAs [17,34]), we discuss how inappropriately enforced/managed PAs affect conclusions. This approach provides the opportunity to discuss effective PA implementations under stochastic population dynamics.

## 2 Methods

In the model, the focal region has two categories: PAs and non-PAs (Fig. 2). I examine the CV in PAs, non-PA, and the whole region for various sizes of PAs, different levels of quality (measured by the degree of decline of mortality rate therein), and several parameters such as site preference, and the degree of stochasticity. The immigration and emigration of individuals to and from PAs connect these areas. The sizes and site preferences affect the likelihood of individual migration. Inhomogeneous mortality rates and sizes in the two regions induce different growth rates *r*_*i*_ and carrying capacities *K*_*i*_, which characterize the *quality of PAs*—higher-quality PAs offer lower mortality in the region, leading to a larger conservation effect. First, I discuss a simple situation in which the population size remains fixed, and introduce some key aspects of the model and analytical results. Then, I discuss a more general situation in which demographic stochasticity occurs in birth, death, and migration events. I also make the explicit connection to existing deterministic models, and thereby introduce SDEs with environmental stochasticity.

**Figure 2:**
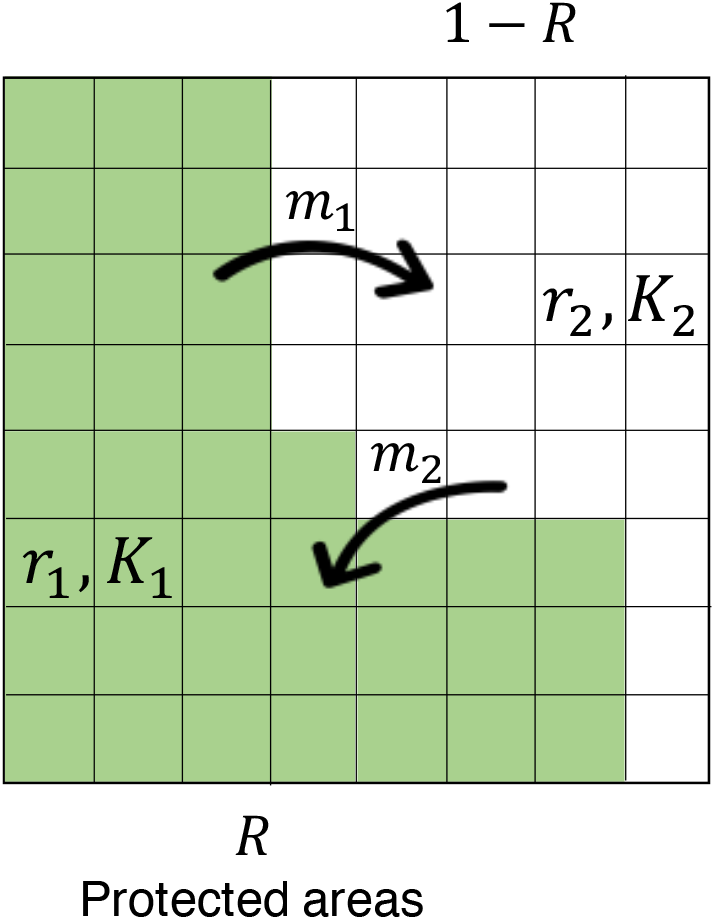
The model scheme. The concerned region is subdivided by subdivisions and these have two categories: PA (fraction *R*; green region) and non-PA (fraction 1-*R*; white region). The emigration from PAs and the immigration to PAs exchange individuals between the areas at constant rates of *m*_1_ and *m*_2_, respectively. Each area is characterized by an intrinsic growth rate *r*_*i*_ and a carrying capacity *K*_*i*_ (*i* = 1, 2), and site preference affects the likelihood of individual migration (see the main text).

When PAs have higher growth rates than non-PAs, it is possible to swap the definition of the two areas. That is, one can regard areas having lower growth rates as PAs, and discuss the effect of “inappropriate” PAs. In the following, I will provide results for both PAs and non-PAs, and the discussion of “good” PAs encompasses “inappropriate” PAs.

### 2.1 Population dynamics with fixed population sizes

I begin with a simple situation in which the population dynamics of a focal species are driven by the immigration and emigration of individuals to and from PAs, and no birth or death events occur. Each area has a site preference, which affects the realized migration rate. The realized migration rate determines the degree of mixture between PAs and non-PAs. Detailed technical discussions can be found in Appendix A.1. The number of individuals remains fixed at *N*, and I write the population sizes in PAs and non-PAs as *n* and *N*− *n*, respectively. Let *p*(*n, t*) be the probability of *n* individuals located in PAs at time *t*. Then, the population dynamics can be described by a simple gain–loss process (Appendix A.1):

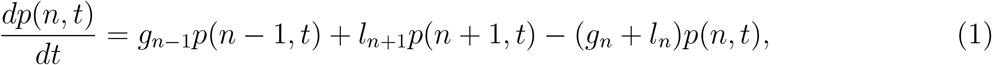

where *g*_*n*_ and *l*_*n*_ are the gain and loss rates corresponding to one individual gain and loss in PAs, respectively. Let *N*_*s*_ and *n*_*s*_ be the total number of subdivisions of the concerned region and the number of subdivisions categorized as PAs, respectively, and let *R* = *n*_*s*_*/N*_*s*_ be the fraction of PAs in the region of interest (Fig. 2). Only the immigration and emigration of individuals drive population changes in PAs, and the gain and loss terms become

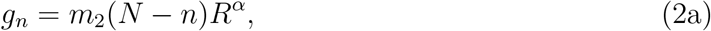

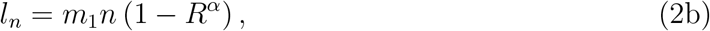

where *m*_1_ and *m*_2_ are the emigration and immigration rates, respectively. The parameter *α* controls the preference of PAs, accounting for preferred/non-preferred/neutral sites by the species (Fig. A.1). Neutral preference (*α* = 1) means that the destination of an individual on the move is determined at random and weighted by the sizes of the PAs and non-PAs. When PAs are preferred (*α <* 1) or non-PAs are preferred (*α >* 1) by the species, the probability of choosing PAs is higher or lower than the neutral choice, respectively.

### 2.2 Population dynamics under demographic stochasticity

When birth and death occur in a population, the total population number *N* is no longer constant, but is the sum of the population sizes in PAs *n*_1_ and non-PAs *n*_2_: *N* = *n*_1_ + *n*_2_. When the population size changes dynamically as a result of births, deaths, and migrations, Eq. (1) becomes (see Appendix A.2 for full details)

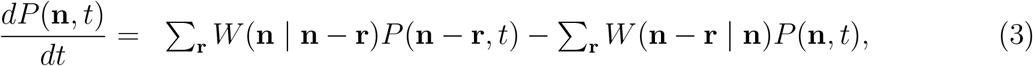

where **r** is a state and *W* (**n | m**) is the transition rate from state **m** to state **n**. These transition rates are

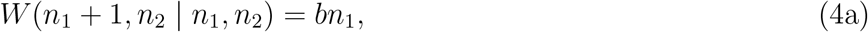

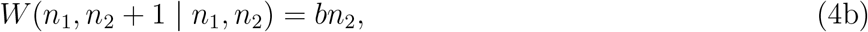

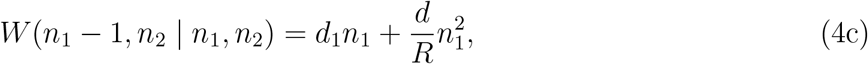

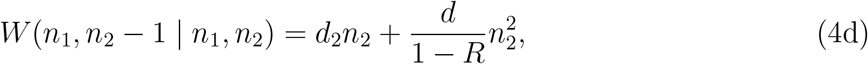

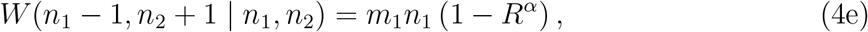

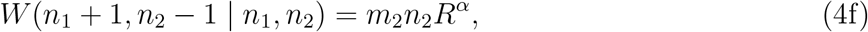

where *b* is the birth rate, *d*_*i*_ is the density-independent mortality rate in site *i* (*i* = 1, 2), and *d* is the density-dependent mortality rate. The factors *R* and 1 − *R* in Eqs. (4c) and (4d) denote the fractions of PAs and non-PAs, respectively. For example, this accounts for a larger density-dependent mortality in a smaller region. The intrinsic growth rate *r*_*i*_ and the carrying capacity *K*_*i*_ in site *i* (*i* = 1, 2) are defined as *r*_*i*_ = *b*− *d*_*i*_ and *K*_*i*_ = *r*_*i*_*/d*, respectively (see Appendix A.2 for details).

The connections to existing deterministic models can be observed by deriving a deterministic representation of Eq. (5). Multiplying both sides by *n*_1_ and summing over all probabilities, I recover a deterministic two-patch dynamics equation (Appendix A.2):

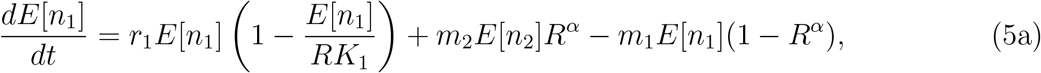

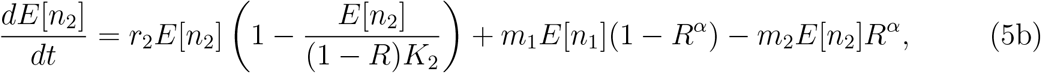

where *E*[*x*] represents the average of *x*. This model is discussed, for example, in [40, 41] with some arrangements.

### 2.3 Population dynamics under environmental stochasticity

I next introduce a stochastic model incorporating environmental stochasticity. This can be obtained from the representation of SDEs [24, 43]. Under the deterministic dynamics of Eq. (5) and with continuous variables *X*_*i*_ representing the population size in PAs (*i* = 1) and non-PAs (*i* = 2), the SDEs can be written as [13]

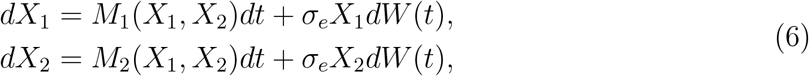

where *M*_*i*_ corresponds to the deterministic part of the population dynamics in region *i* (Eq. A.19) and *σ*_*e*_ is the intensity of environmental stochasticity.

## 3 Results

### 3.1 Long-term effects of PAs with a fixed population size

When population changes in PAs are solely due to migrations between PAs and non-PAs, I can obtain analytical insights that are not possible for more general situations. Later, I will show that analytical insights from this simple situation can be applied to these more general situations. In the model, I can derive explicit forms of the mean and the CV in PAs as follows (Appendix A.1):

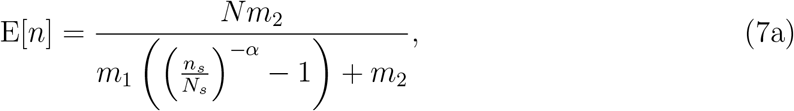

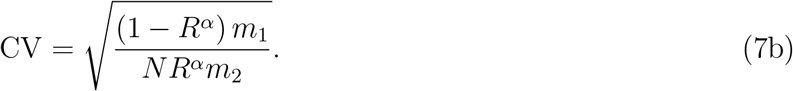

The parameter dependence on the CV in PAs is analyzed by differentiating Eq. (7b) with respect to the parameter of interest. For example, the relationship *∂*CV*/∂m*_2_ *<* 0 indicates that an increase in the immigration rate to PAs *m*_2_ decreases the CV, and *∂*CV*/∂α >* 0 indicates that an increase of the site preference for non-PAs *α* increases the CV, respectively. In addition, I conclude that an increase in the PA fraction always decreases the CV in PAs because *∂*CV*/∂R <* 0. Figure 3 represents the relationships among the expressions in Eqs. (7), and confirms the above discussions.

**Figure 3:**
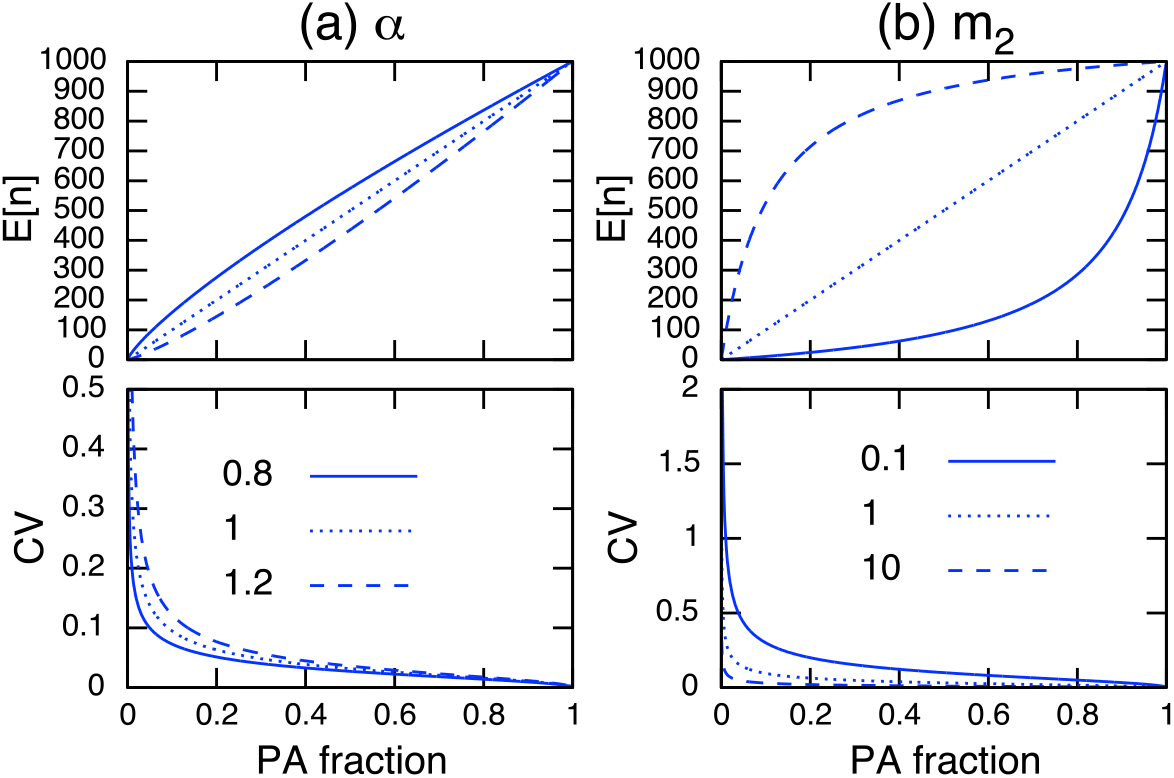
Parameter dependence on E[*n*] (top) and the CV in PAs (bottom) of site preference *α* (left) and of immigration rate to PAs *m*_2_ (right). Other parameter values used are *N* = 1000, *m*_1_ = 1 (left), and *α* = 1 (right). Note the site preference is *α <* 1 when PAs are preferred and *α >* 1 when non-PAs are preferred.

I can further simplify Eq. (7b) by setting E[*n*] = *n* in Eq. (7a), solving for an arbitrary parameter, and plugging the result into Eq. (7b). This cancels *R, α, m*_1_, and *m*_2_ in Eq. (7b), and the CV in PAs can be described only by the population size in PAs, *n*, and the total population size, *N* (Eq. (A.14) in Appendix A.1). This indicates that a larger population size in PAs results in a smaller CV. Hence, I conclude that any parameters that improve the population size in PAs reduce its CV.

### 3.2 Long-term effects of PAs under demographic stochasticity

When birth and death events occur in addition to migrations, the total population size changes dynamically. Although this situation is not amenable to mathematical analysis because of the nonlinearity in the demographic rate terms, numerical simulations imply that I can still discuss this situation following a similar line to the analytical discussions above. For instance, Figs. 4 (a) and (b) show that the site preference *α*, immigration rate to PAs *m*_2_, and PA fraction have a qualitatively similar dependence on the CV in PAs to that of the fixed-population scenario (Fig. 3).

**Figure 4:**
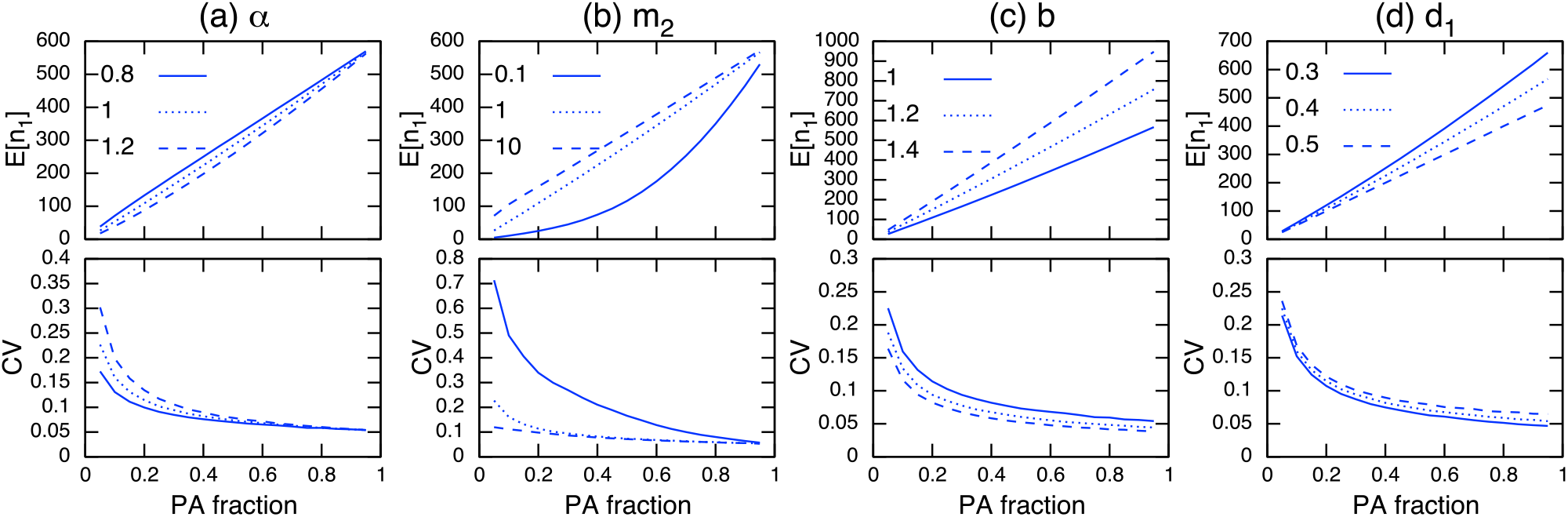
Parameter dependence on E[*n*_1_] (top) and CV in protected areas (bottom) under demographic stochasticity. The parameter being examined is specified at the top of each column. Other parameter values used (unless specified) are *b* = 1, *d* = 0.001, *d*_1_ = 0.4, *d*_2_ = 0.5, *m*_1_ = 1, *m*_2_ = 1, and *α* = 1. Note the site preference is *α <* 1 when PAs are preferred and *α >* 1 when non-PAs are preferred.

Regarding the dependence of demographic parameters, I can use the relationships obtained in the preceding analysis. That is, changes in parameter values that increase the population size in PAs will reduce its CV (Fig. 4c, d). This can be heuristically seen using the result in the preceding analysis (Appendix A.1), whereby a larger population size in PAs, *n*_1_, leads to a smaller CV in PAs, while the magnitude of the population size in non-PAs, *n*_2_, scales with the CV:

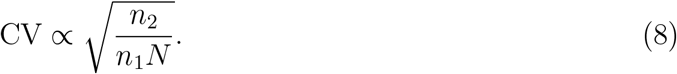

Generally speaking, expanding the PAs increases their population size and decreases that in non-PAs. If a conservation effect of PAs is sufficient, it also improves the total population size *N*. This leads to a smaller CV in PAs as the PA fraction increases (Fig. 4).

Figure A.2 shows the results for non-PAs and the whole region. The opposite trend from that in PAs can be observed: increasing the PA fraction increases the CV in non-PAs, which is associated with a population decline in the region. Additionally, the total population size determines the CV in the whole region.

### 3.3 Long-term effects of PAs under environmental stochasticity

Environmental stochasticity affects the whole population, and its influence does not vanish even with large population sizes, unlike demographic stochasticity. Under environmental stochasticity, the relationships discussed above are no longer useful, and the interpretation of results becomes more intricate. For example, increasing the PA fraction and population size in PAs does not guarantee a reduction in the CV, but can instead increase the CV (Fig. 5b; immigration rates *m*_2_=0.1 and 10). However, these actions may reduce the CV when *m*_2_ = 1.0. Similarly, the site preference *α* exhibits a complex response in terms of the effect on the CV in PAs (Fig. 5a), and for PAs with a higher preference, increasing the PA fraction may increase the CV (Fig. 5a; *α* = 0.8 and PA fraction *R <* 0.2).

**Figure 5:**
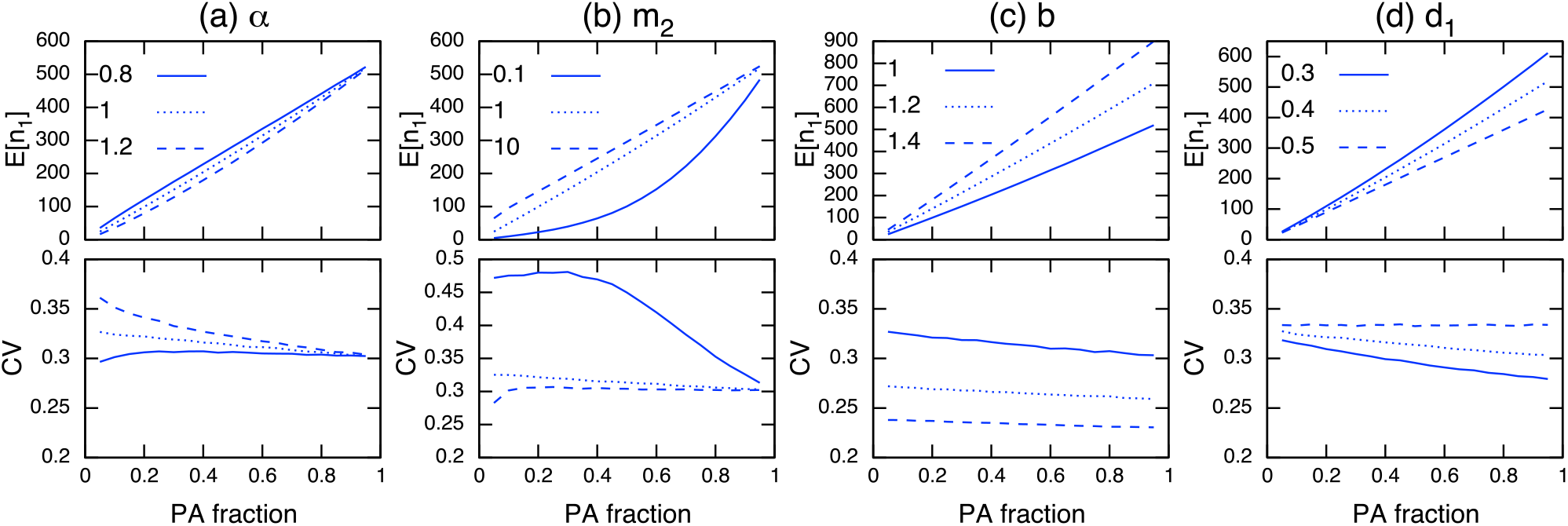
Parameter dependence on E[*n*_1_] (top) and CV in PAs (bottom) under environmental stochasticity. The parameter being examined is specified at the top of each column. Other parameter values used (unless specified) are *b* = 1, *d* = 0.001, *d*_1_ = 0.4, *d*_2_ = 0.5, *m*_1_ = 1, *m*_2_ = 1, *α* = 1, and 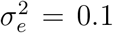 Note the site preference is *α <* 1 when PAs are preferred and *α >* 1 when non-PAs are preferred.

The establishment of PAs is likely to decrease the corresponding CV if a conservation effect of PAs is increased by reducing the density-independent mortality rate in PAs, *d*_1_ (Fig. 5d). Increasing the birth rate *b* (a focal species has a high fecundity rate) shows a similar effect, and tends to decrease the CV regardless of the PA fraction (Fig. 5c). However, a larger birth rate decreases the conservation effects of PAs (e.g., the relative difference between the net growth rate of PAs and non-PAs (*b − d*_1_)*/*(*b* − *d*_2_) decreases when *b* increases), and the effect of reducing the CV in PAs becomes smaller. These numerical observations can be verified analytically when the immigration and emigration rates are sufficiently large (*m*_1_, *m*_2_ ≫ 1). In this condition, the CV in PAs is described as (Appendix A.3)

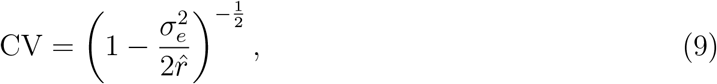

where 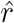 can be interpreted as an average population growth rate in the whole region weighted by the site preference (*α*). From Eq. (9), I conclude that an increase in the PA fraction decreases the CV in PAs if the growth rate in PAs is greater than that in non-PAs (*b* − *d*_1_ *> b*− *d*_2_), and *vice versa*. I also state that a small PA size has a large effect on reducing the CV when the site preference for PAs is higher than that for non-PAs (*α <* 1), and *vice versa* (Appendix A.3).

Results for non-PAs and the whole region are provided in Fig. A.3, but the CV trends in these cases are not easy to distinguish from those in PAs. This highlights the difference from the situations under demographic stochasticity (Figs. 4 and A.2; bottom), where the opposite trends in population sizes and CVs occur between the two areas. This may be because environmental stochasticity affects all individuals regardless of the population size, and the stochastic influence tends to be consistent across the whole region. Additionally, the CV in the whole area is more likely to decrease with increasing total population size (Fig. A.3). However, this is not the case when the immigration rate is large (Fig. A.3f; *m*_2_ = 10).

When PAs provide a higher conservation effect (*d*_1_ = 0.1), the trend for an increasing CV in PAs would be mitigated (Fig. A.4b; *m* = 0.1, 10), or even reversed (Fig. A.4a; *α* = 0.8). I also examined a situation with a higher environmental stochasticity of 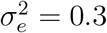, and obtained qualitatively similar results (Fig. A.5).

## 4 Discussion

### General findings

A multitude of indices can characterize the effect of PAs [25]. Here, I have analyzed the long-term conservation effects of PAs under stochasticity, which is not simply an equilibrium discussion. I measured the long-term PA effects via the CV of the population size, where PAs with a small CV offer less variation, and hence have a larger long-term conservation effect. Loosely speaking, the CV in PAs is suppressed by increasing their area if the PAs provide a sufficient conservation effect. Hence, the implementation of effective PAs promotes long-term conservation effects. This general finding agrees with previous studies focused on the marine environment [5, 20, 31].

### Effects of demographic/environmental stochasticity

Demographic stochasticity is suppressed by a large population size [24]. Therefore, if PAs increase the population size by, for example, expanding in size or increasing the immigration probability (Figs. 3, 4a, b), or reducing the mortality rate (Fig. 4d), the population fluctuations are suppressed; this is a mechanism for achieving long-term conservation effects. Moreover, if PAs increase the total population size in the focal region, fluctuations in the population are suppressed by the same mechanism (Fig. A.2e–h). The developed master equation approach allows us to derive a concrete analytical insight for a fixed population size, e.g., under the assumption that the target species has a low growth rate compared to the timescale of a conservation program. In fact, many conservation practices act over much shorter timescales than ecological timescales [11, 48].

In contrast, environmental stochasticity affects all individuals, and population size plays a minor role in determining the long-term conservation effects (Fig. 5a, b, and d). However, if the conservation effect of PAs is high (e.g., mortality is sufficiently reduced in PAs), expanding the PAs tends to reduce the CV. Equivalently, the CV in PAs is increased by introducing ineffective PAs (e.g., mortality is not sufficiently reduced or increased in PAs). In addition, if PAs increase the total population size in the whole region, the CV tends to decrease in that region (Fig. A.3), except for situations with a high immigration rate, where the opposite trend is observed (e.g., Fig. A.3f; *m*_2_ = 10).

### Inappropriate PAs may amplify fluctuations

In the proposed model, the density-independent mortality rate characterizes the quality of PAs, and I implicitly assumed that PAs reduced the mortality rate in those areas. In fact, this definition is arbitrary, and one can argue that PAs have a higher mortality than non-PAs, a situation that could be described as “inappropriate” PAs. The results of this case are already discussed as “non-PAs” in this paper (e.g., Fig. A.2a–d). Alternatively, an increase in mortality inside PAs would reverse patterns identified here (i.e., proceeding right to left along *x*-axes in Figs. 3-5): increasing PA fraction would increase overall mortality, thereby increasing the CV.

In practice, poorly implemented PAs can arise due to inappropriate planning or management process, such as lack of sufficient commutations between stakeholders [2]. Existing unfairness affects compliance of management and can increase conflict between user groups [2]. Illegal use of protected species is one of the potential risks of PAs management that can increase the mortality of a target species [34]. For instance, Harasti et al. [17] reported that illegal fishing activities potentially reduced the abundance of a fish species by 55% from 2011-2017 in the Seal Rocks no-take area in Australia. Similarly, Hopf et al. [20] demonstrated certain PAs designs can further destabilize a system. Hence, effective enforcement of PAs is necessary to promote its benefit.

### Strategies to achieve robust PA management

I have identified different mechanisms for enhancing the population size in PAs, such as those to improve the demographic rate (i.e., birth *b* and death *d*_1_) and to promote the probability of remaining in PAs (i.e., site preference *α*, emigration *m*_1_ and immigration *m*_2_ rates). In practice, if PAs adequately regulate anthropogenic activities and are monitored (e.g., strict nature reserve [22]), the demographic rate may be improved in PAs, providing a long-term conservation effect under demographic/environmental stochasticity. However, ineffective PA management is often associated with a failure to reduce human activities [8,37], which may amplify population fluctuations. The immigration and emigration rates are not purely biological parameters, but can be controlled by the configuration of the PAs [40]. Protecting the favored sites of a target species is desirable as a means of increasing the conservation effects of PAs [21]. However, when MPAs are used as a sustainable fisheries management tool, a moderate spillover effect is necessary to promote fishing yields [29, 38]. Under environmental stochasticity, this can lead to a reduced long-term effect of PAs, and careful assessment is necessary.

MPAs can improve the resilience of populations under existing multiple stable states [1, 5, 42]. The findings in this paper further complement this insight. Namely, while a catastrophic shift of population is incurred by population perturbations [36], adequately established PAs tend to suppress population fluctuations (i.e., the population is more robust against perturbations). Therefore, populations within PAs are more likely to remain in the basin of attraction of a current stable state. A more explicit discussion addressing the relationships between the CV of the population dynamics, the strength of perturbations, and the basin of attraction will further improve our understanding.

Likewise, there are multiple directions for further extending this analysis. For example, here, I have assumed that environmental stochasticity affects PAs and non-PAs equally. While this is reasonable when the concerned region is small, and the environment in each type of area is not different, two areas may be subject to other environmental fluctuations [16]. In fact, the size of an MPA sometimes becomes significant [10], and the model developed needs to incorporate heterogeneous environmental stochasticities. Age and metapopulation structures have been used in the context of fisheries management [20], and these investigations are also relevant to the context of this paper. While the study discussed the expected population size along with the CV provided by a fraction of PAs established, an expected timescale to observe such a population size is an important management consideration [4,23]. Kaplan et al. [23] demonstrated recovery is fast at small population sizes where variations of population fluctuations are also small. Revealing the role of the CV in population recovery will further complement the current knowledge. However, it should be noted that applying complex models has a large cost in terms of reduced generality and analytical intractability. With this in mind, the approach of the study will guide the development of a general framework to discuss the long-term conservation effects of PAs.

## Funding

The University of Tokyo provided funding for this project. Also, this work was partially supported by JSPS KAKENHI Grant Number 21K17913.

## Acknowledgements

I am grateful to T. Fung and Steven D. Aird for their comments on the manuscript. I am also thankful to two anonymous reviewers for their valuable comments.

## Data Availability Statement

No data were collected for this study.

## Conflict of Interest

I declare I have no conflict of interest.

## Appendix Figures

**Figure A.1:**
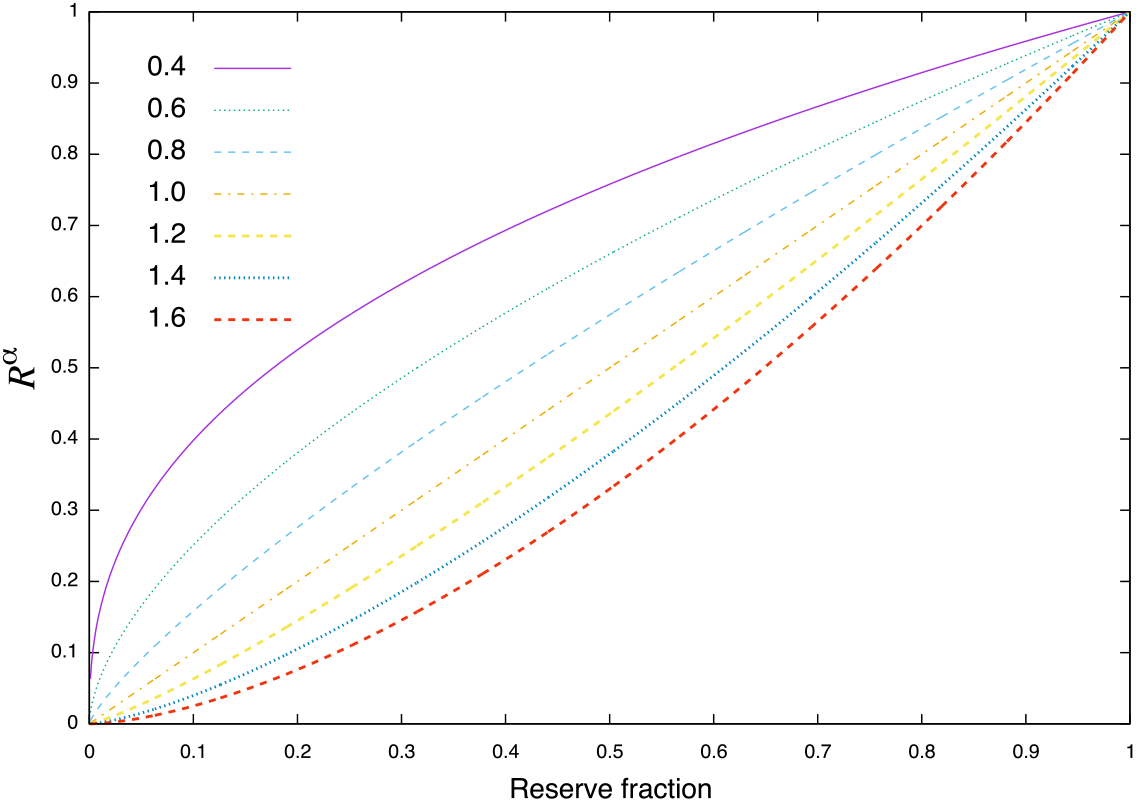
Effect of the site preference *α* on the probability of immigration to PAs given the fraction of PAs *R*. When *α* = 1, migration occurs at random.

**Figure A.2:**
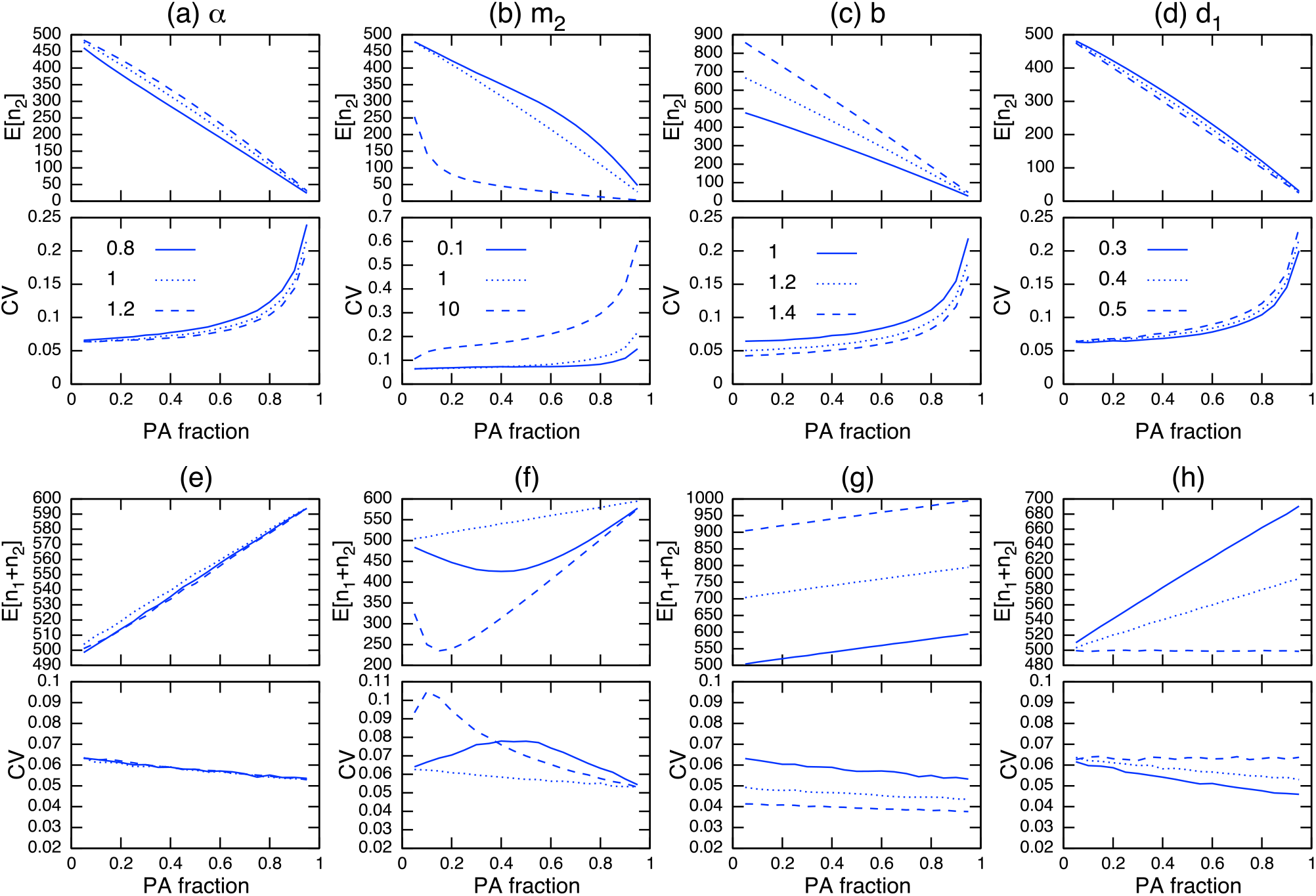
Parameter dependence on E[*n*_2_] and CV of *n*_2_ (a)–(d); and E[*n*_1_ + *n*_2_] and CV of *n*_1_ + *n*_2_ (e)–(h) under demographic stochasticity. The parameter being examined is specified at the top of each column. Other parameter values used (unless specified) are *b* = 1, *d* = 0.001, *d*_1_ = 0.4, *d*_2_ = 0.5, *m*_1_ = 1, *m*_2_ = 1, and *α* = 1.

**Figure A.3:**
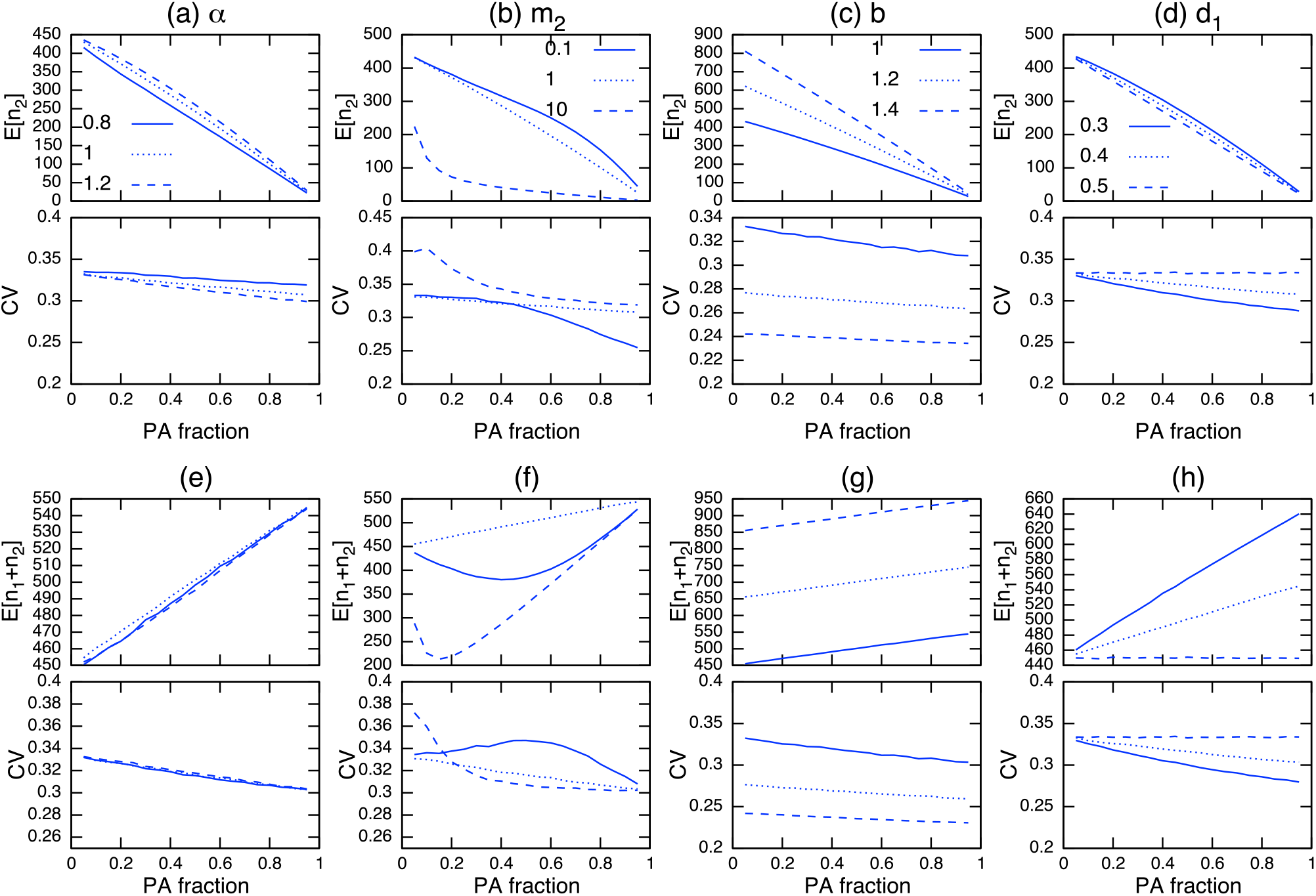
Parameter dependence on E[*n*_2_] and CV of *n*_2_ (a)–(d); and E[*n*_1_ + *n*_2_] and CV of *n*_1_ + *n*_2_ (e)–(h) under environmental stochasticity with intensity 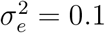 The parameter being examined is specified at the top of each column. Other parameter values used (unless specified) are *b* = 1, *d* = 0.001, *d*_1_ = 0.4, *d*_2_ = 0.5, *m*_1_ = 1, *m*_2_ = 1, and *α* = 1.

**Figure A.4:**
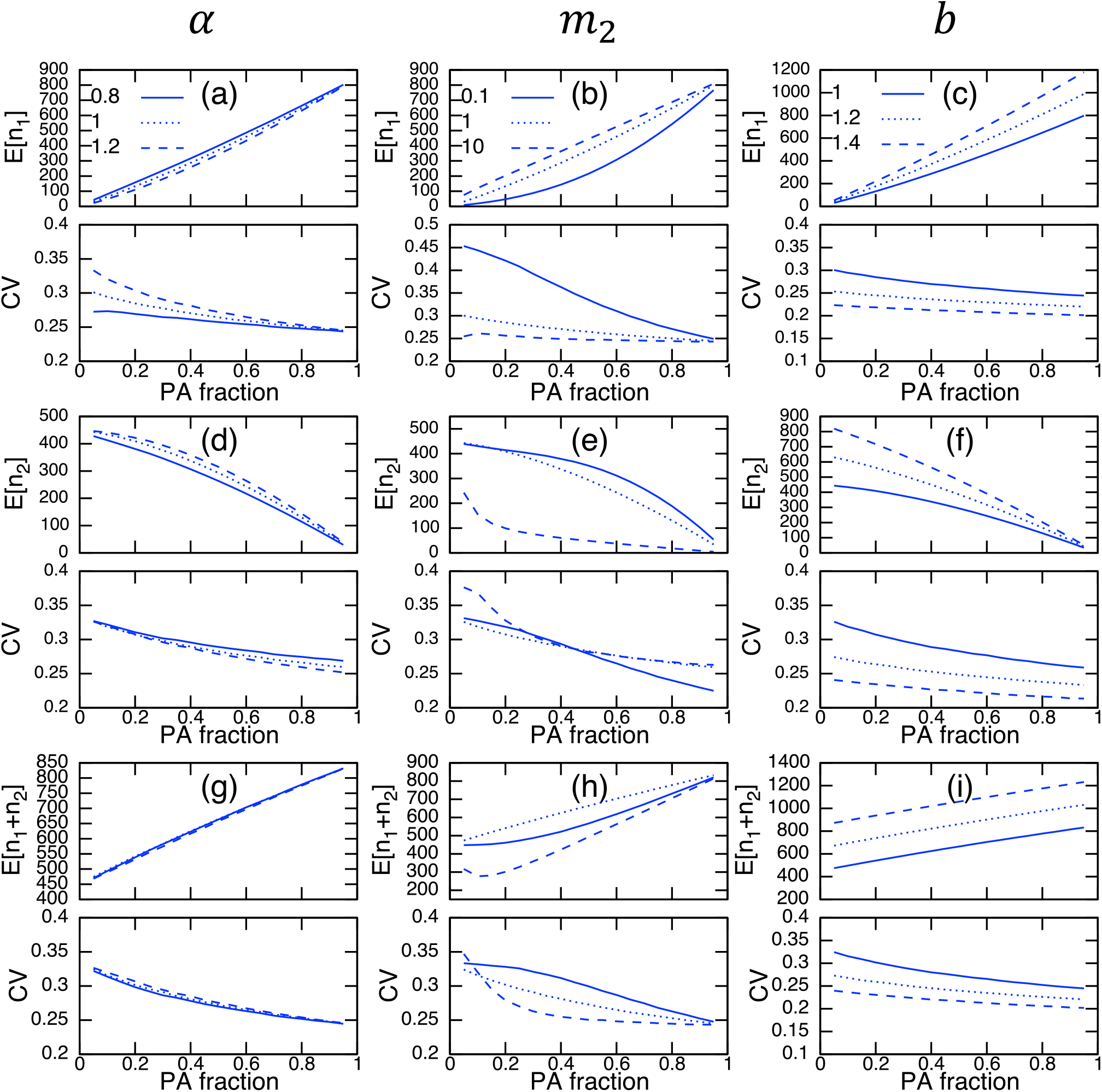
Parameter dependence on E[*n*_1_] and CV of *n*_1_ (a)–(c); E[*n*_2_] and CV of *n*_2_ (d)–(f); and E[*n*_1_ + *n*_2_] and CV of *n*_1_ + *n*_2_ (g)–(i) under environmental stochasticity with intensity 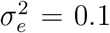 The parameter being examined is specified at the top of each column. Other parameter values used (unless specified) are *b* = 1, *d* = 0.001, *d*_1_ = 0.1, *d*_2_ = 0.5, *m*_1_ = 1, *m*_2_ = 1, and *α* = 1.

**Figure A.5:**
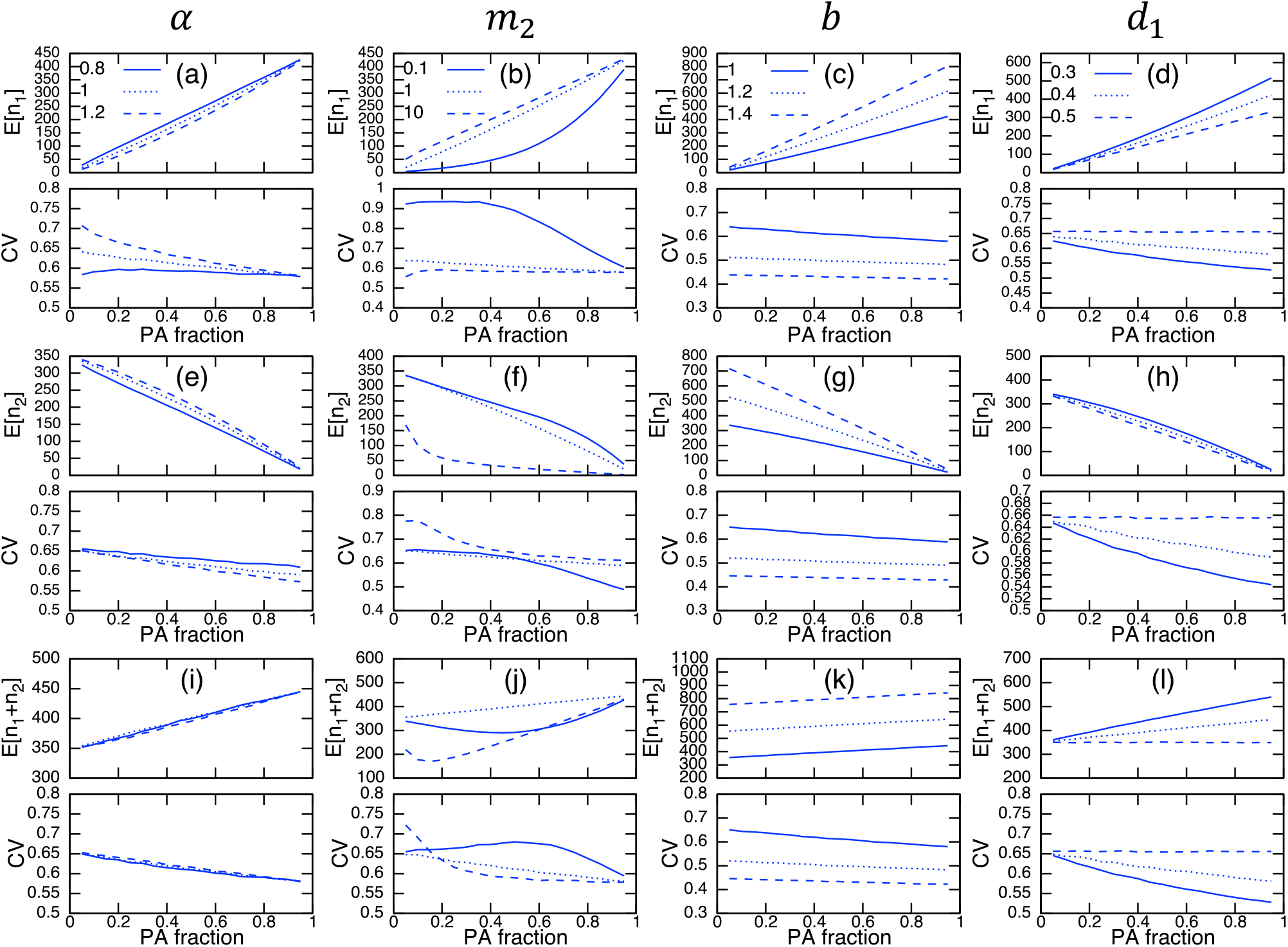
Parameter dependence on E[*n*_1_] and CV of *n*_1_ (a)–(d); E[*n*_2_] and CV of *n*_2_ (e)–(h); and E[*n*_1_ + *n*_2_] and CV of *n*_1_ + *n*_2_ (i)–(l) under environmental stochasticity with intensity 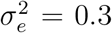. The parameter being examined is specified at the top of each column. Other parameter values used (unless specified) are *b* = 1, *d* = 0.001, *d*_1_ = 0.4, *d*_2_ = 0.5, *m*_1_ = 1, *m*_2_ = 1, and *α* = 1.

**Figure A.6:**
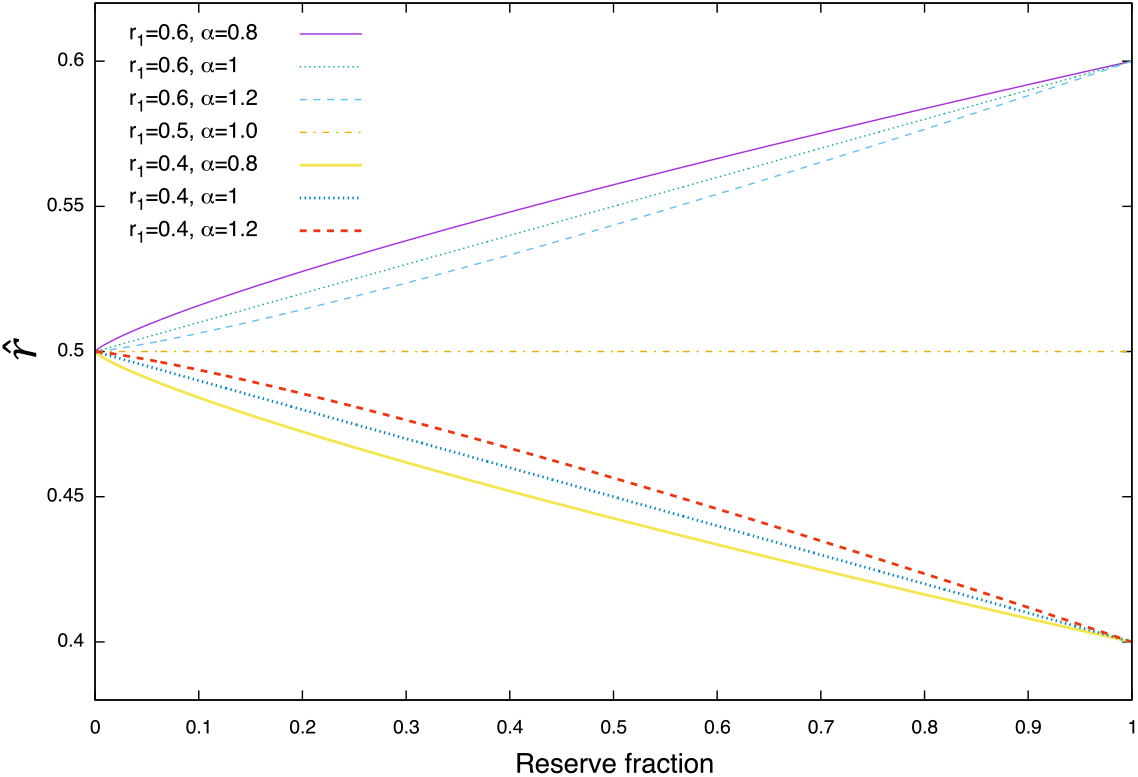
Parameter dependence on 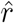 (Eq. A.23a); *r*_2_ = 0.5.

## Appendix

### A.1 Population dynamics with fixed population size

I begin with a simple situation in which the population dynamics of a focal species are driven by the immigration and emigration of individuals to and from protected areas (PAs), and not by birth and death events. Each area has a site preference, which affects the realized migration rate between the two areas. Let *p*(*n, t*) be the probability of *n* individuals located in PAs at time *t*. Then, the population change is described by the following relationship:

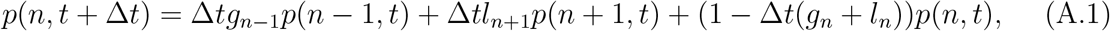

where *g*_*n*_ and *l*_*n*_ are the gain and loss probabilities corresponding to a gain and a loss of an individual in PAs. Equation (A.1) leads to the following gain–loss process:

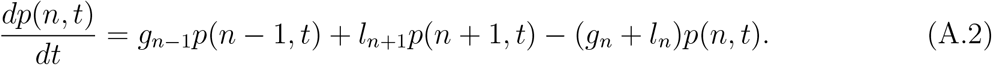

Let *N*_*s*_ and *n*_*s*_ be the total number of subdivisions of the concerned region and number of subdivisions categorized as PAs, respectively, and define the PA fraction as *R* = *n*_*s*_*/N*_*s*_. Then, individual gains and losses are the result of immigration and emigration, and the gain and loss terms become

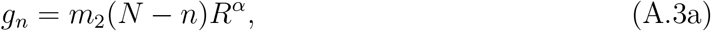

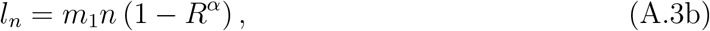

where *m*_1_ and *m*_2_ are the emigration and immigration rates, respectively. The parameter *α* controls the preference for PAs. From Eq. (A.3b), it can be seen that *l*_0_ = 0. By setting *dp/dt* = 0 in Eq. (A.2), I get a stationary state

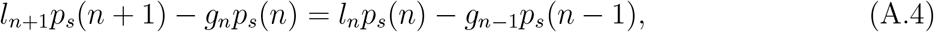

which is satisfied by all *n* = 0, 1, *…*. By setting *n* = 0, I get

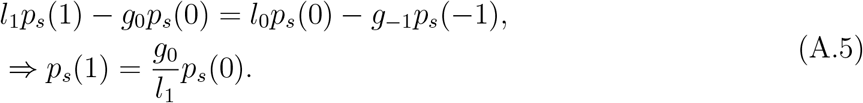

Therefore, I have

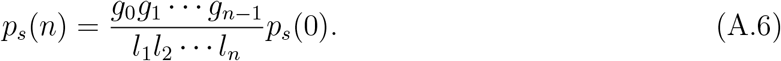

The probability should be normalized to 1, and *p*(0) here is the normalizing factor. This is calculated as

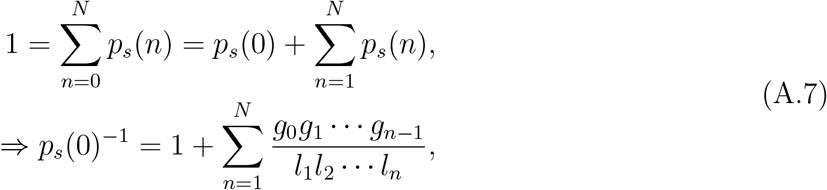

where, with the two expressions in Eq. (A.3), the last expression becomes

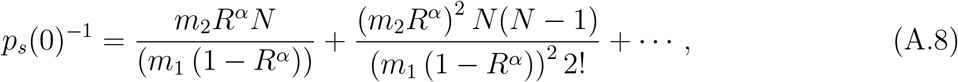

and its *n*th term is described as

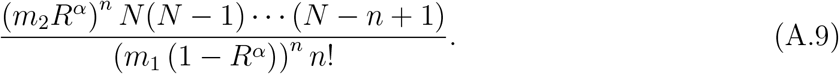

This expression leads to

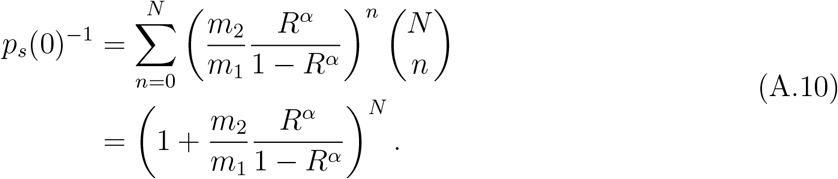

Hence, substituting this into Eq. (A.6), I obtain the stationary probability

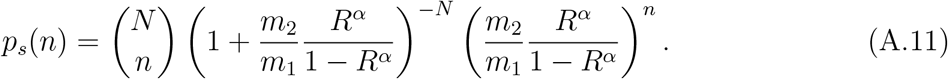

The mean, variance, and coefficient of variation (CV) are calculated from Eq. (A.11) as

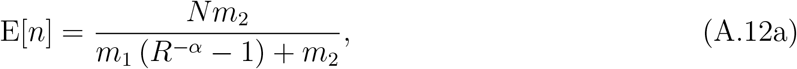

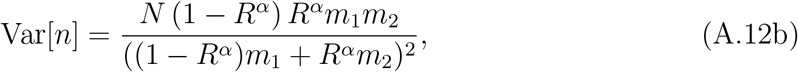

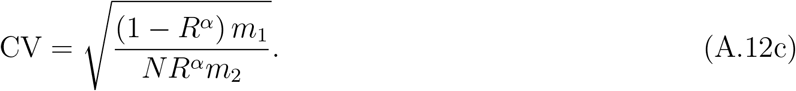

The parameter dependence on the CV can be determined by differentiating Eq. (A.12c) with respect to the parameter of interest:

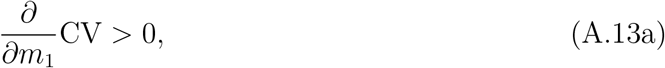

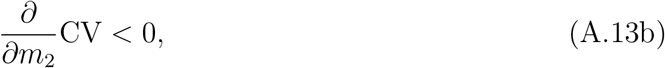

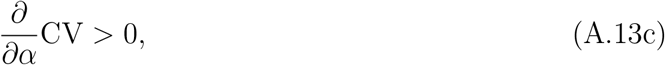

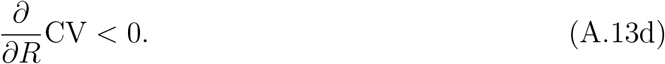

Furthermore, by setting E[*n*] = *n* in Eq. (A.12a) and solving for an arbitrary parameter, and then plugging the result into Eq. (A.12c), I have the following relationship:

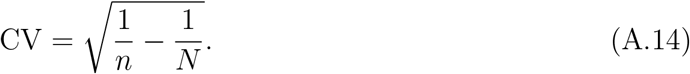

Hence, the CV is a function of *n* and *N* only, and the parameters in the model affect the PA effect by changing the population size *n* in the PAs.

### A.2 Population dynamics under demographic stochasticity

Here, I extend the gain–loss process of Eq. (A.2) to incorporate population changes caused by birth and death events. Let *n*_1_ and *n*_2_ be the population sizes in PAs and non-PAs, respectively, and *d*_1_ and *d*_2_ be the density-independent mortality rates in PAs and non-PAs, respectively. When the total population size *N* changes dynamically, the gain–loss process of Eq. (A.2) becomes

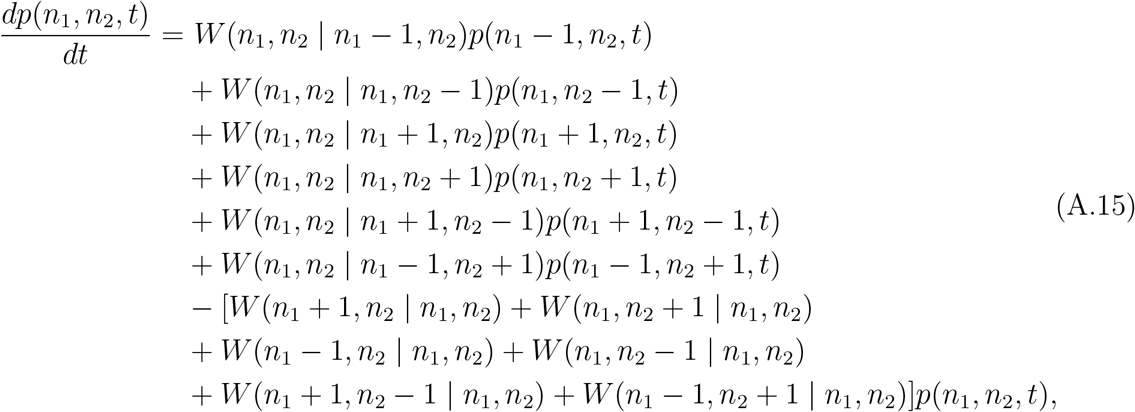

where 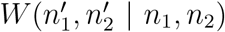 is the transition probability from state (*n*_1_, *n*_2_) to 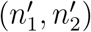. These transition probabilities are

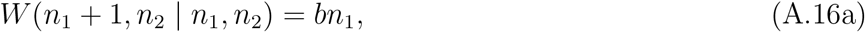

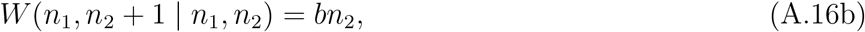

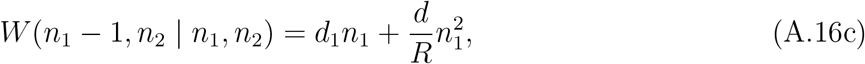

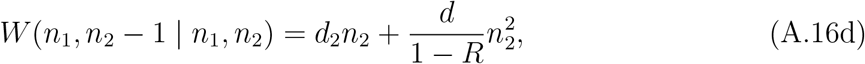

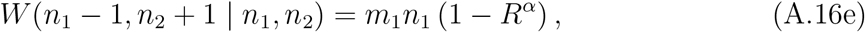

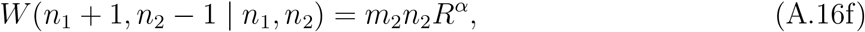

where *R* and 1 − *R* in Eqs. (A.16c) and (A.16d) account for the size of the PAs and non-PAs, respectively, in the density-dependent mortality rate *d*, so as to induce a larger density-dependent mortality in a smaller region. This model is not amenable to analytical approaches because of the nonlinearity in the transition probabilities. Hence, I need to rely on numerical simulations. However, I heuristically found that the CV in PAs is scaled by the relationship in Eq. (A.14), but the total population size *N* is also a random variable in this situation:

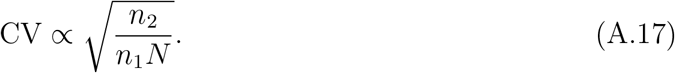

The deterministic dynamics of *n*_1_ from Eq. (A.19) are derived by multiplying by *n*_1_ on both sides and summing over all possible probabilities. This gives

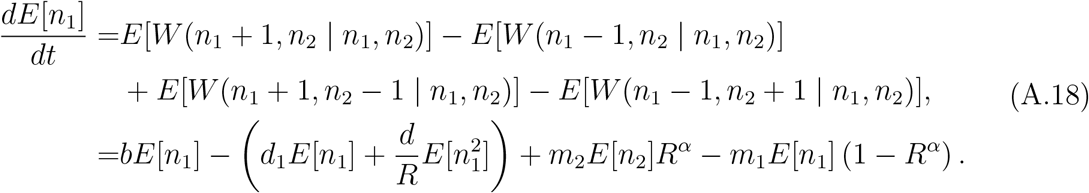

With the decoupling approximation 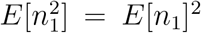, and together with the dynamics of *E*[*n*_2_], I have

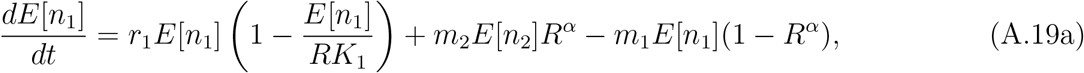

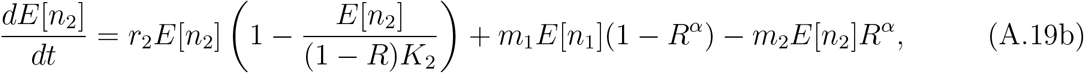

where *r*_*i*_ = *b* − *d*_*i*_ and *K*_*i*_ = *r*_*i*_*/d* are the growth rate and carrying capacity in area *i*.

### A.3 Aggregated model and its stochastic dynamics

When the emigration and immigration rates are large (*m*_1_, *m*_2_ ≫ 1), I can aggregate the deterministic dynamics of Eq. (A.19). Let *x*_1_ and *x*_2_ be the population sizes, denoted by continuous variables, in PAs and non-PAs, respectively, and *X* = *x*_1_ + *x*_2_ be the total population size. Then, I write the aggregate dynamics as [1,2]

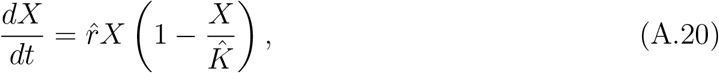

where, at equilibrium, I have

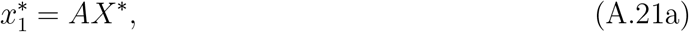

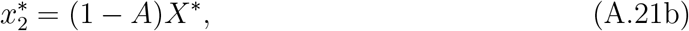

in which 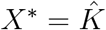 and *A* = *m*_2_*R*^*α*^*/*(*m*_1_(1 − *R*^*α*^) + *m*_2_*R*^*α*^). The aggregated parameters 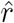 and 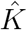 are defined as

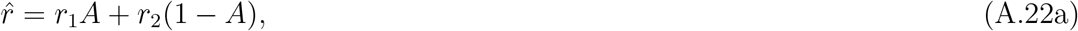

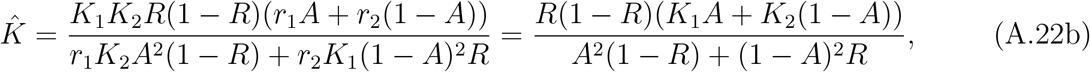

where *r*_*i*_ = *b*_*i*_ − *d*_*i*_ and *K*_*i*_ = (*b*_*i*_ − *d*_*i*_)*/d* are the growth rate and carrying capacity in area *i*. At the limit as *m*_1_, *m*_2_ → ∞, I have 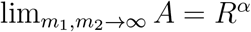, and the expressions in Eq. (A.22) become

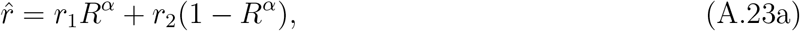

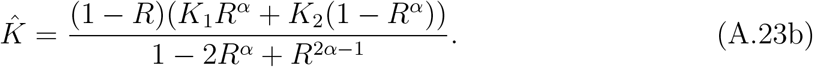

Alternatively, setting *m*_2_ = *Bm*_1_ and taking the limit as *m*_1_ → ∞, I have

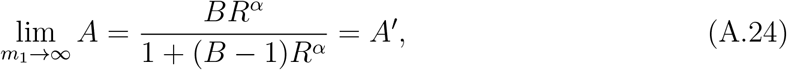

where the coefficient *B* represents the ratio of *m*_1_ and *m*_2_, and *B* = 1 realizes *A* = *R*^*α*^. Substituting this expression into Eq. (A.22) allows the asymmetric migrations between PAs and non-PAs to be examined. The stochastic differential equation can be described using the notation of Eq. (A.20):

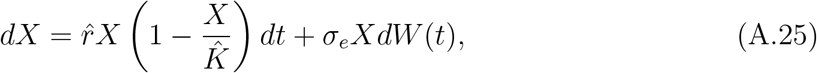

where *X* is now a random variable. In Itô’s interpretation, this is equivalent to the following Fokker–Planck equation [3]:

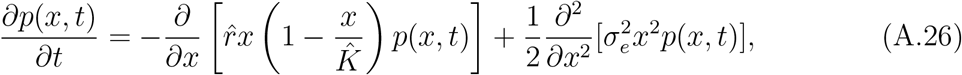

which has a quasi-stationary distribution of the form of a Gaussian distribution:

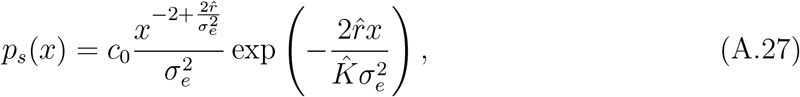

where 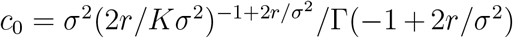 is a normalization constant. This expression gives the mean, variance, and CV of *x* as follows:

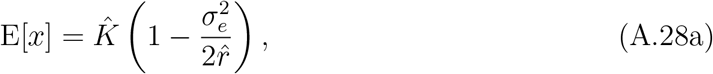

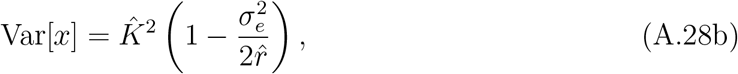

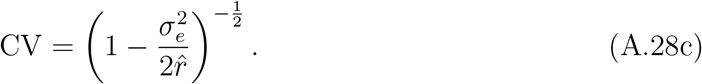

By taking the derivative of the CV (Eq. A.28c) with respect 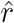 under the positivity to requirement 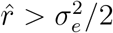, I have

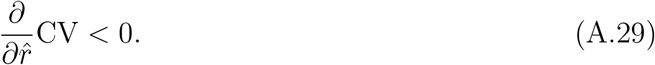

This means that an increment in 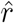 reduces the CV. From Eq. (A.22a), I see that the value 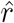 can be increased by increasing the PA fraction *R* when *r*_1_ *> r*_2_. That is, if PAs have a larger intrinsic growth rate than non-PAs, introducing such “good” PAs reduces the CV. The parameter *α* controls the sensitivity of this effect; *α <* 1 gives the largest rate of reduction of the CV (i.e., a curve with diminishing returns) when *R* is small, and the opposite trend is realized when *α >* 1. These are depicted in Fig. A.6. Note that Eq. (A.27) is the probability in the whole region concerned, but the carrying capacity 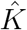 is canceled in the expression of the CV (Eq. A.28c). This also suggests that the subregions (i.e., PAs and non-PAs) of the concerned region have the same CV in the large-value limit of *m*_1_ and *m*_2_, as expected.

## Notes

### Competing Interest Statement

The authors have declared no competing interest.

## References

[1] E. A. Aalto, F. Micheli, C. A. Boch, J. A. Espinoza Montes, C. B. Woodson, and G. A. De Leo. Catastrophic mortality, allee effects, and marine protected areas. m. Nat., 193:391–408, 2019.

[2] T. Agardy, G. N. di Sciara, and P. Christie. Mind the gap: Addressing the shortcomings of marine protected areas through large scale marine spatial planning. Mar. Policy, 35:226–232, 2011.

[3] C. M. Aiken and S. A. Navarrete. Environmental fluctuations and asymmetrical dispersal: Generalized stability theory for studying metapopulation persistence and marine protected areas. Mar. Ecol. Prog. Ser., 428:77–88, 2011.

[4] C. Barceló, J. W. White, L. W. Botsford, and A. Hastings. Projecting the timescale of initial increase in fishery yield after implementation of marine protected areas. ICES J. Mar. Sci., page fsaa233, 2021.

[5] L. A. Barnett and M. L. Baskett. Marine reserves can enhance ecological resilience. Ecol. Lett., 18:1301–1310, 2015.

[6] W. C. Clark. Mathematical bioeconomics: the optimal management of renewable resources. John Wiley & Sons, Inc, New York, 2nd edition, 1990.

[7] C. Costello and S. Polasky. Optimal harvesting of stochastic spatial resources. J. Environ. Econ. Manage., 56:1–18, 2008.

[8] I. D. Craigie, J. E. Baillie, A. Balmford, C. Carbone, B. Collen, R. E. Green, and J. M. Hutton. Large mammal population declines in Africa’s protected areas. Biol. Conserv., 143:2221–2228, 2010.

[9] G. A. De Leo and F. Micheli. The good, the bad and the ugly of marine reserves for fishery yields. Philos. Trans. R. Soc. B Biol. Sci., 370:20140276, 2015.

[10] N. K. Dulvy. Super-sized MPAs and the marginalization of species conservation. Aquat. Conserv. Mar. Freshw. Ecosyst., 23:357–362, 2013.

[11] C. A. Froyd and K. J. Willis. Emerging issues in biodiversity & conservation management: The need for a palaeoecological perspective. Quat. Sci. Rev., 27:1723–1732, 2008.

[12] J. M. Fryxell, D. H. Lynn, and P. J. Chris. Harvest reserves reduce extinction risk in an experimental microcosm. Ecol. Lett., 9:1025–1031, 2006.

[13] C. Gardiner. Stochastic Methods - A Handbook for the Natural and Social Sciences. Springer-Verlag, Berlin, 2009.

[14] J. Geldmann, M. Barnes, L. Coad, I. D. Craigie, M. Hockings, and N. D. Burgess. Effectiveness of terrestrial protected areas in reducing habitat loss and population declines. Biol. Conserv., 161:230–238, 2013.

[15] R. Q. Grafton, T. Kompas, and D. Lindenmayer. Marine reserves with ecological uncertainty. Bull. Math. Biol., 67:957–971, 2005.

[16] H. Hakoyama and Y. Iwasa. Extinction risk of a meta-population: Aggregation approach. J. Theor. Biol., 232:203–216, 2005.

[17] D. Harasti, T. R. Davis, A. Jordan, L. Erskine, and N. Moltschaniwskyj. Illegal recreational fishing causes a decline in a fishery targeted species (Snapper: Chrysophrys auratus) within a remote no-take marine protected area. PLoS One, 14:e0209926, 2019.

[18] A. Hastings, S. D. Gaines, and C. Costello. Marine reserves solve an important bycatch problem in fisheries. Proc. Natl. Acad. Sci. U. S. A., 114:8927–8934, 2017.

[19] M. H. Holden, D. Biggs, H. Brink, P. Bal, J. Rhodes, and E. McDonald-Madden. Increase anti-poaching law-enforcement or reduce demand for wildlife products? A frame-work to guide strategic conservation investments. Conserv. Lett., 12:e12618, 2018.

[20] J. K. Hopf, G. P. Jones, D. H. Williamson, and S. R. Connolly. Marine reserves stabilize fish populations and fisheries yields in disturbed coral reef systems. Ecol. Appl., 29:e01905, 2019.

[21] T. N. Hunt, S. J. Allen, L. Bejder, and G. J. Parra. Identifying priority habitat for conservation and management of Australian humpback dolphins within a marine protected area. Sci. Rep., 10:14366, 2020.

[22] IUCN. Guidelines for applying protected area management categories. 2008.

[23] K. A. Kaplan, L. Yamane, L. W. Botsford, M. L. Baskett, A. Hastings, S. Worden, and J. W. White. Setting expected timelines of fished population recovery for the adaptive management of a marine protected area network. Ecol. Appl., 29:e01949, 2019.

[24] R. Lande, S. Engen, and B.-E. Saether. Stochastic Population Dynamics in Ecology and Conservation. Oxford University Press, Oxford, 2003.

[25] F. Leverington, K. L. Costa, H. Pavese, A. Lisle, and M. Hockings. A global analysis of protected area management effectiveness. Environ. Manage., 46:685–698, 2010.

[26] M. Mangel. Irreducible uncertainties, sustainable fisheries and marine reserves. Evol. Ecol. Res., 2:547–557, 2000.

[27] M. Mangel. On the fraction of habitat allocated to marine reserves. Ecol. Lett., 3:15–22, 2000.

[28] C. R. Margules and R. L. Pressey. Systematic conservation planning. Nature, 405:243– 53, 2000.

[29] T. R. McClanahan and S. Mangi. Spillover of exploitable fishes from a marine park and its effect on the adjacent fishery. Ecol. Appl., 10:1792–1805, 2000.

[30] A. J. McKane and T. J. Newman. Stochastic models in population biology and their deterministic analogs. Phys. Rev. E, 70:041902, 2004.

[31] C. Mellin, M. Aaron Macneil, A. J. Cheal, M. J. Emslie, and M. Julian Caley. Marine protected areas increase resilience among coral reef communities. Ecol. Lett., 19:629–637, 2016.

[32] R. M. Nisbet and W. Gurney. Modelling Fluctuating Populations. Wiley, New York, 1982.

[33] H. Possingham, I. Ball, and S. Andelman. Mathematical methods for identifying representative reserve networks. In S. Ferson and M. Burgman, editors, Quant. methods Conserv. Biol., pages 291–305. Springer-Verlag New York, New York, USA, 2000.

[34] J. H. Razafimanahaka, R. K. Jenkins, D. Andriafidison, F. Randrianandrianina, V. Rakotomboavonjy, A. Keane, and J. P. Jones. Novel approach for quantifying illegal bushmeat consumption reveals high consumption of protected species in Madagascar. ORYX, 46:584–592, 2012.

[35] E. Sala and S. Giakoumi. No-take marine reserves are the most effective protected areas in the ocean. ICES J. Mar. Sci., 75:1166–1168, 2018.

[36] M. Scheffer, S. Carpenter, J. Foley, and C. Folke. Catastrophic shifts in ecosystems. Nature, 413:591–596, 2001.

[37] K. Schulze, K. Knights, L. Coad, J. Geldmann, F. Leverington, A. Eassom, M. Marr, S. H. Butchart, M. Hockings, and N. D. Burgess. An assessment of threats to terrestrial protected areas. Conserv. Lett., 11:e12435, 2018.

[38] B. Stobart, R. Warwick, C. González, S. Mallol, D. Díaz, O. Reñones, and R. Goñi. Long-term and spillover effects of a marine protected area on an exploited fish community. Mar. Ecol. Prog. Ser., 384:47–60, 2009.

[39] N. Takashina. Simple rules for establishment of effective marine protected areas in an age-structured metapopulation. J. Theor. Biol., 391:88–94, 2016.

[40] N. Takashina. On the spillover effect and optimal size of marine reserves for sustainable fishing yields. PeerJ, 8:e9798, 2020.

[41] N. Takashina, J.-H. Lee, and H. Possingham. Effect of marine reserve establishment on non-cooperative fisheries management. Ecol. Modell., 360:336–342, 2017.

[42] N. Takashina and A. Mougi. Effects of marine protected areas on overfished fishing stocks with multiple stable states. J. Theor. Biol., 341:64–70, 2014.

[43] M. Turelli. Random environments and stochastic calculus. Theor. Popul. Biol., 12:140–178, 1977.

[44] O. Venter, R. A. Fuller, D. B. Segan, J. Carwardine, T. Brooks, S. H. Butchart, M. Di Marco, T. Iwamura, L. Joseph, D. O’Grady, H. P. Possingham, C. Rondinini, R. J. Smith, M. Venter, and J. E. Watson. Targeting Global Protected Area Expansion for Imperiled Biodiversity. PLoS Biol., 12:e1001891, 2014.

[45] J. E. Watson, N. Dudley, D. B. Segan, and M. Hockings. The performance and potential of protected areas. Nature, 515:67–73, 2014.

[46] C. D. West, C. Dytham, D. Righton, and J. W. Pitchford. Preventing overexploitation of migratory fish stocks: The efficacy of marine protected areas in a stochastic environment. ICES J. Mar. Sci., 66:1919–1930, 2009.

[47] E. R. White, M. L. Baskett, and A. Hastings. Catastrophes, connectivity, and Allee effects in the design of marine reserve networks. Oikos, 2020.

[48] K. J. Willis, L. Gillson, T. M. Brncic, and B. L. Figueroa-Rangel. Providing baselines for biodiversity measurement. Trends Ecol. Evol., 20:107–108, 2005.

## References

[1] Y. Iwasa, V. Andreasen, and S. Levin. Aggregation in model ecosystems. I. Perfect aggregation. Ecol. Model., 37:287-302, 1987.

[2] N. Takashina. On the spillover effect and optimal size of marine reserves for sustainable fishing yields. PeerJ, 8:e9798, 2020.

[3] C. Gardiner. Stochastic Methods - A Handbook for the Natural and Social Sciences. Springer-Verlag, Berlin, 2009.

